# *In situ* administration of cytokine combinations induces tumor regression

**DOI:** 10.1101/260547

**Authors:** Jinyu Zhang, Haochen Jiang, Yunhai Zhang

**Affiliations:** Mianyi Biotechnology Corporation, Chongqing 401332, China.; Center for Life Sciences, Tsinghua University, Beijing 100084, China.; Beijing Chaoyang District Animal Disease Control Center, Beijing 100018, China.

## Abstract

Recent advances in cancer immunotherapy suggest a possibility of harnessing the immune system to defeat malignant tumors, but complex immunosuppressive microenvironment confined the therapeutic benefits to a minority of patients with solid tumors. Here we constructed a lentivector based inducible system to evaluate the therapeutic effect of cytokines in established tumors. Doxycycline (Dox) induced local expression of cytokine combinations exhibited strong synergistic effect, leading to complete regression of tumors. Notably, IL12+GMCSF+IL2 expression induced eradication of tumors in all mice tolerated with this treatment, including those bearing large tumors of ~15mm in diameter, and generated an intensive systemic antitumor immunity. Other combinations with similar immune regulatory roles also induced tumor elimination in a majority of mice. Moreover, intratumoral injection of chitosan/IL12+GMCSF+IL2 solution induced complete response in all tested syngeneic tumor models, regardless of various tumor immunograms. These results provide a versatile method for the immunotherapy of intractable malignant neoplasm.

## Main Text

In recent years, cancer immunotherapy, including immune checkpoint inhibition, CAR-T, et al, has become a powerful weapon for fighting malignant neoplasm (1–8). The results from experimental mouse tumor models and human clinic trails indicate that the immune system has the potential of eradicating malignant tumors, even those in advanced stage. However, the clinic outcome is unsatisfactory, e.g., the low response rate of immune checkpoint therapeutics and the ineffectiveness of CAR-T in solid tumors (9–11). There remains a need for new modalities that available to harness the immune system to defeat solid tumors. Here we report a finding that in situ administration of cytokines is able to efficiently eliminate the primary tumors in experimental mice.

Inducible expression systems have been used to evaluate the influences of some genes or siRNAs on tumor development (12–15). We constructed a lentivector based doxycycline (dox)- inducible system comprising of two components: 1, a lentiviral vector for constitutive expression of reverse tet transactivator (rtTA); 2, a lentiviral vector harboring a tet-responsive TRE promoter and a MCS site, into which cytokine genes are easy to be cloned (Fig. 1A). After transduction and antibiotics selection, B16F10 cells carrying various inducible genes were generated. Addition of dox induced a significant expression of cloned gene in vitro (fig. S1). Using iRFP as a marker (16), we tested the availability of this system in vivo. Fluorescence could be detected 24h post induction, and gradually increased in the following days (Fig. 1B). Notably, survival analysis suggested that the tumorigenicity of B16F10 cells was not altered by lentiviral transduction and dox administration (fig. S2). Based on that, the therapeutic effects of some cytokines, which has the capability of immune activation, were assessed with this system. Though been approved in cancer therapy in clinic (17), monotherapy of some cytokines could only suppressed tumor growth, instead of eliminating established tumors in this system. IL12, IFNα and IFNγ induced tumor regression in part of tested mice (Fig. 1C, fig. S3). Subsequently, the synergistic antitumor effects of cytokine combinations were investigated. In the antitumor immunity, macrophages and T cells played pivotal roles, as well as dendritic cells are important for the tumor antigen presentation. Three cytokines, IL12, IL2 and GMCSF, capable of activating these cells, were selected to evaluate combination effects. All mice in IL12+IL2 group, 86.7% in IL12+GMCSF group and 40% in GMCSF+IL2 group were cured, which indicated a strong synergistic effect (Fig. 1D). Furthermore, the combination of IL12, GMCSF and IL2 exhibited superior antitumor activity, that all tumors were eliminated within 10 days after induction (Fig. 1E). The ulceration of tumor area suggested an existence of acute immune attack (Fig. 1F). Considering the importance of IFNγ in antitumor immunity, we additionally tested the curative effects of IL12+GMCSF+IL2+IFNγ combination. Interestingly, the addition of IFNγ attenuated the antitumor activity of IL12+GMCSF+IL2 combination, leading to a delayed tumor regression (Fig. 1G). It suggested that an elaborate design of cytokine combination was required for maximal stimulation of antitumor immunity.

**Fig. 1.**
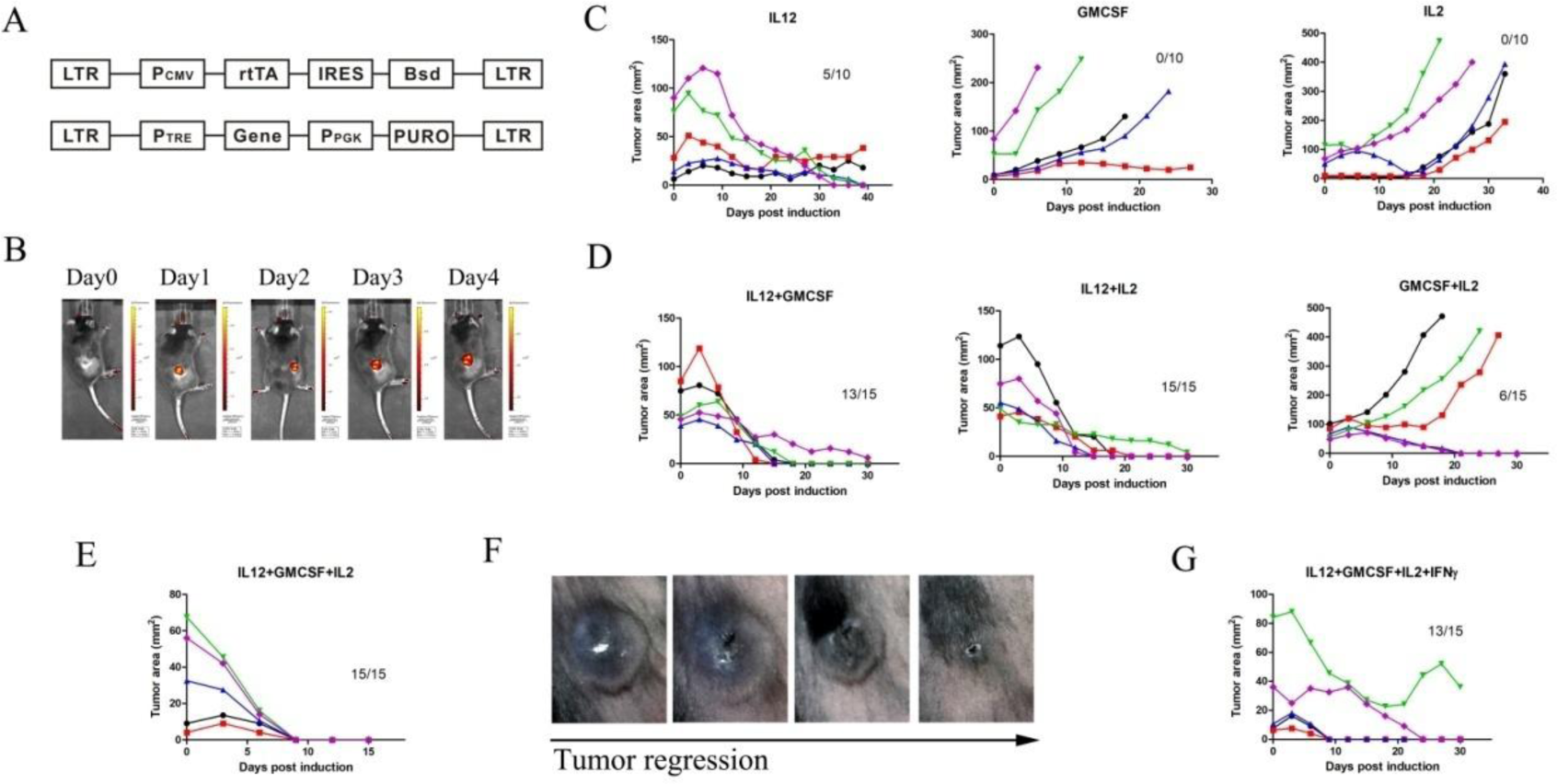
A lentivector based inducible system for evaluating the antitumor effects of cytokines. (A) Schematic representation of the two lentivectors of the inducible system. (B) Dox-induced iRFP expression in B16F10-rtTA(TRE-iRFP) tumors. (C) Effects of dox-induced single cytokine expression, including IL12, GMCSF or IL2, on the growth of established tumors. (D) Effects of dox-induced two-cytokine combinations expression, including IL12+GMCSF, IL12+IL2 or GMCSF+IL2, on the growth of established tumors. (E) Effects of dox-induced expression of IL12+GMCSF+IL2 combination on the growth of established tumors. (F) Representative photographs of tumor regression after induced IL12+GMCSF+IL2 expression. (G) Effects of dox-induced expression of IL12+GMCSF+IL2+IFN γ combination on the growth of established tumors. Each curve represented the tumor growth of an individual mouse after dox administration. Numbers in the graphs indicated tumor cleared mice/total dox treated mice.

Next, we assessed the therapeutic potential of IL12+GMCSF+IL2 combination on late stage melanoma. Addition of dox induced tumor cells to secret high level cytokines (fig. S4), which provided a sustained cytokine supply in the inducible tumors. For advanced tumors (longest diameter range from 10mm to 15mm), induction of cytokine expression led to a complete clearance in all tumor bearing mice (Fig. 2A). In the group of mice with larger tumor burden (longest diameter over 15mm), dox addition induced elimination of tumors in all tolerated mice (Fig. 2A, fig. S5). Of note, except for curative, the other mice died within 4 days after induction, which was likely due in part to an overwhelming inflammation from immune attack to large tumors. Then we rechallenged the IL12+GMCSF+IL2 cured mice, subcutaneously or intravenously, with parental B16F10 cells over 2 month later. All mice rejected inoculated tumor cells, suggesting an immune memory targeting endogenous tumor antigens (Fig. 2B). To investigate whether the immune response could erase established tumors, parental tumor cells were inoculated contralaterally to inducible tumors. Induction of cytokine expression significantly prolonged the survival time of mice bearing a small parental tumor and a large inducible tumor (Fig. 2C, fig. S6A). In contrast, antitumor immunity raised from cytokine expression in small inducible tumors had little effect on the growth of large parental tumors on contralateral flank (Fig. 2C, fig. S6B). However, if the parental and inducible tumors were both in a larger size, all mice died within 6 days after induction (Fig. 2C, fig. S6C). Thereafter, we evaluated the effects of antitumor immune response on metastatic tumor cells. Parental B16F10 cells were intravenously injected into mice bearing inducible tumors. Long term survival was achieved in all injected mice if dox was administered just after injection (Fig. 2D, fig. S7A). If dox was given 2 days post injection, 40% mice cleared intravenous tumor cells and the remaining died within 4 days after induction (Fig. 2D, fig. S7B). It was likely due to a systemic inflammation raised from immune response targeting visceral metastatic lesions. Unlike other immunotherapy modality (18, 19), the side effect after cytokine treatment was much slighter, as vitiligo was well restricted at tumor site (Fig. 2E).

**Fig. 2.**
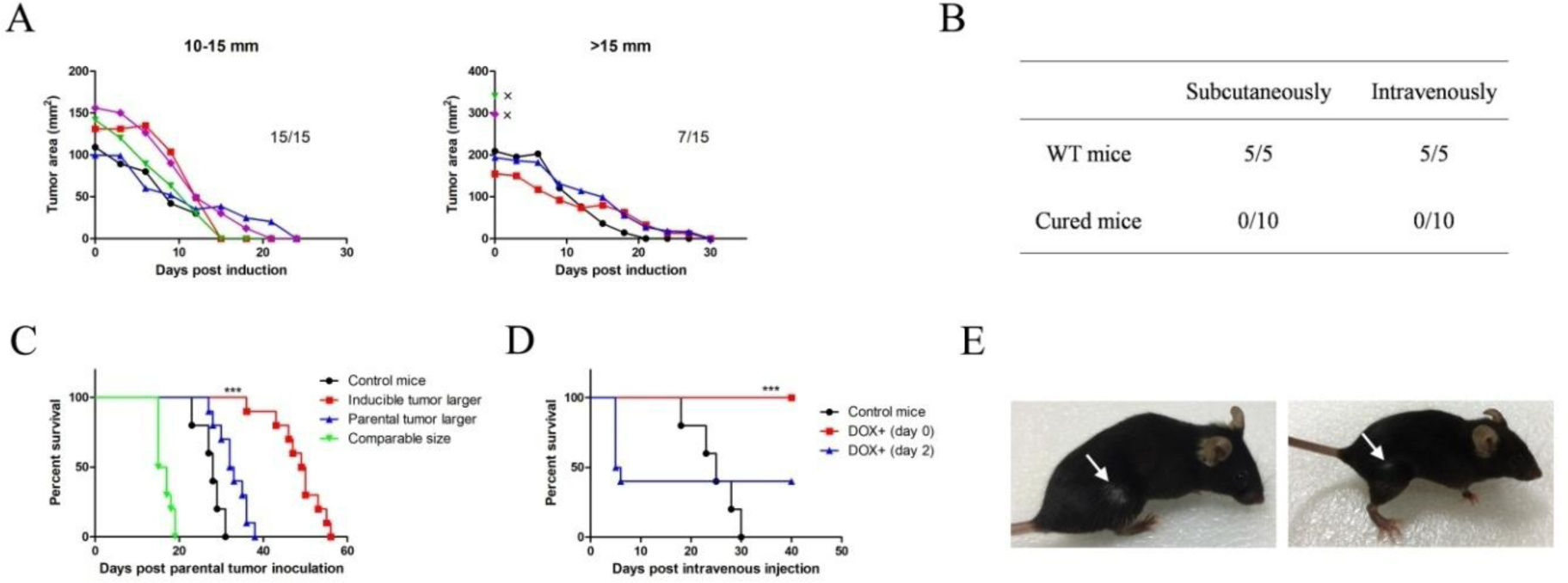
Induction of IL12+GMCSF+IL2 combination expression elicited systemic antitumor immunity. (A) Effects of dox-induced expression of IL12+GMCSF+IL2 combination on the growth of large established tumors (10mm<diameter<15mm or diameter>15mm). Each curve represented the tumor growth of an individual mouse after dox administration. Numbers in the graphs indicated tumor cleared mice/total dox treated mice. X indicated a death case within 4 days after induction. (B) Cured mice rejected rechallenged B16F10 tumor cells, either subcutaneously or intravenously injected. Numbers indicated death case/injected mice. (C) At different times post subcutaneous inoculation of inducible tumor cells into the flank of C57BL/6 mice, parental tumor cells were subcutaneously injected into the contralateral side. Dox induction was initiated when the size of bilateral tumors at different ratio (inducible tumor larger, parental tumor larger, or size of both tumors were comparable). Mice bearing only parental tumors served as control. Survival of tumor bearing mice was recorded. n=5 for control group and n=10 for other groups. ***, *p*<0.001. (D) Parental tumor cells were intravenously injected into mice bearing subcutaneously established inducible tumors. Dox induction was initiated just after injection or 2 days later. Mice without inducible tumors served as control. Survival of tumor bearing mice was recorded. n=5 for control group and n=10 for other groups. ***, *p* <0.001. (E) Vitiligo at tumor site after tumor regression caused by induced expression of IL12+GMCSF+IL2.

To find out cell types participating in the tumor clearance mediated by IL12+GMCSF+IL2 expression, tumor mass were collected and subjected to IHC analysis. Dox induction did not alter the immune cell infiltration in parental tumors. Before induction, there were a little more F4/80+, CD11c+ and CD3+ cells in the tissue of inducible tumors than that in parental tumors, which might be due to the background expression of cytokines (fig. S4). After induction, large amount of macrophages and T cells infiltration was observed in tumors, accompanied with increased number of dendritic cells (fig. S8). This is consistent with previous conclusion that T cells are the predominant effector cells in cancer immunotherapy (20, 21).

Cytokines are multifunctional biomolecules in the immune system, which meant that some cytokines had overlapped activities. Like GMCSF, FLT3L promoted the activation and antigen presentation of dendritic cells. Besides IL2, other cytokines, including IL15 and IL21, had the capacity of promoting T cell activation and expansion (22). Therefore, substituting GMCSF with FLT3L, and IL2 with IL15 or IL21, we tested the antitumor activities of other 5 cytokine combinations: IL12+GMCSF+IL15 (Fig. 3A), IL12+GMCSF+IL21 (Fig. 3B), IL12+FLT3L+IL15 (Fig. 3C), IL12+FLT3L+IL21 (Fig. 3D) and IL12+FLT3L+IL2 (Fig. 3E). All combinations exhibited strong antitumor activities. Specifically, background secretion of IL12+FLT3L+IL2 displayed great suppressive effect in the process of tumor engraftment, as only 50% mice inoculated with these tumor cells generated a palpable tumor lesion. Though tumor growth was markedly inhibited in these groups, only IL12+GMCSF+IL21 induced a complete regression in all mice. Generally, compared to FLT3L, GMCSF exhibited a higher efficiency of stimulating immune clearance of established tumors. Unlike IL12+GMCSF+IL2, vitiligo was rarely observed in cured mice of these groups (fig. S9). The IL12+GMCSF+IL2 induced tumor regression process was also faster than that in these groups (Fig. 1E). Taken together, local administration of cytokine combination by induced expression has the capability of eradicating large established tumors with advanced malignancy, and a better therapeutic effect is in usual accompanied with a higher side effect.

**Fig. 3.**
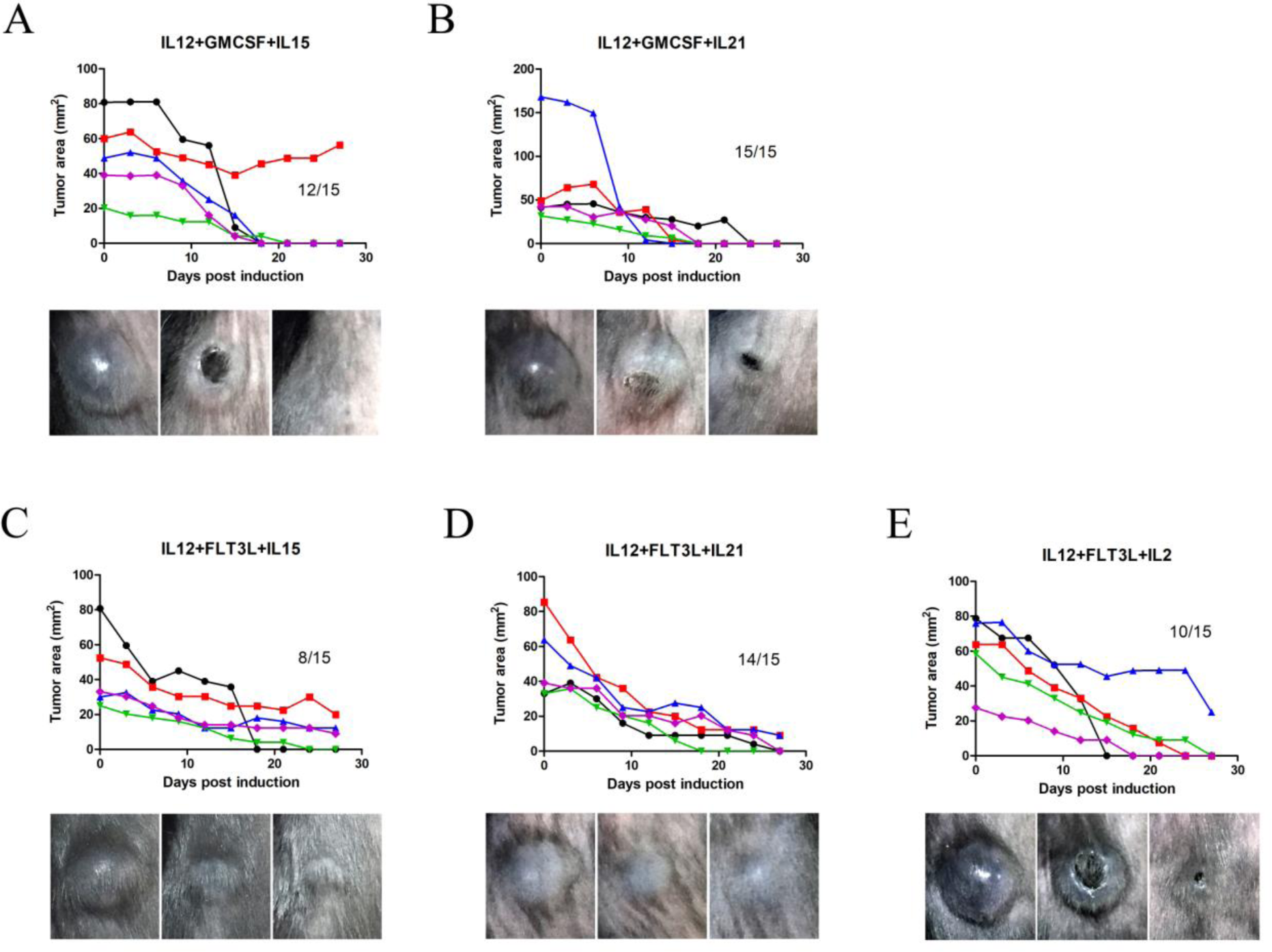
Induction of cytokine combinations expression eliminated established tumors. Mixture of different cytokine inducible B16F10 cells, including IL12+GMCSF+IL15 (A), IL12+GMCSF+IL21 (B), IL12+FLT3L+IL15 (C), IL12+FLT3L+IL21 (D), IL 12+FLT3L+IL2 (E), were subcutaneously inoculated at the flank of C57BL/6 mice. Dox induction was initiated after tumor nodule formation. Growth of tumors was recorded after induction. Each curve represented the tumor growth of an individual mouse after dox administration. Numbers in the graphs indicated tumor cleared mice/total dox treated mice. Photographs indicated a representative tumor regression progress.

Induction of cytokine expression from tumor cells provided a sustained and concentrated cytokine supplement in tumor microenvironment, which is important for persistent immune stimulation and reduced peripheral toxicity. Some slow release materials, e.g., alginate microsphere and chitosan, have been used in cancer therapy to recapitulate this effect (23, 24). Firstly, we constructed 293 cells expressing cytokines by lentiviral transduction (fig. S10A). Using alginate gel as a carrier, cells expressing IL12, GMCSF and IL2 were injected into B16F10 melanoma lesions. After 3-4 rounds of injection, 70% mice successfully cleared established tumors (fig. S11). To further improve the versatility of cytokine treatment, IL12, GMCSF as well as IL2 expressed from 293 cells were concentrated by ultrafiltration (fig. S10B). Mixed with chitosan, the concentrated cytokines were injected into tumors for a long-term retention. Compared to cytokine expressing cells, this treatment was more efficient as one or two injections induced B16F10 melanoma regression in all mice (Fig. 4A). Although there were relapses in part of mice, secondary injection made a regression again. It indicated that there was no resistance to this therapeutics in recurrent tumors. In spite of high antitumor activity in the process of tumor engraftment, delivery of IL12+FLT3L+IL2 with chitosan was not able to induce elimination of established B16F10 tumors (fig. S12).

**Fig. 4.**
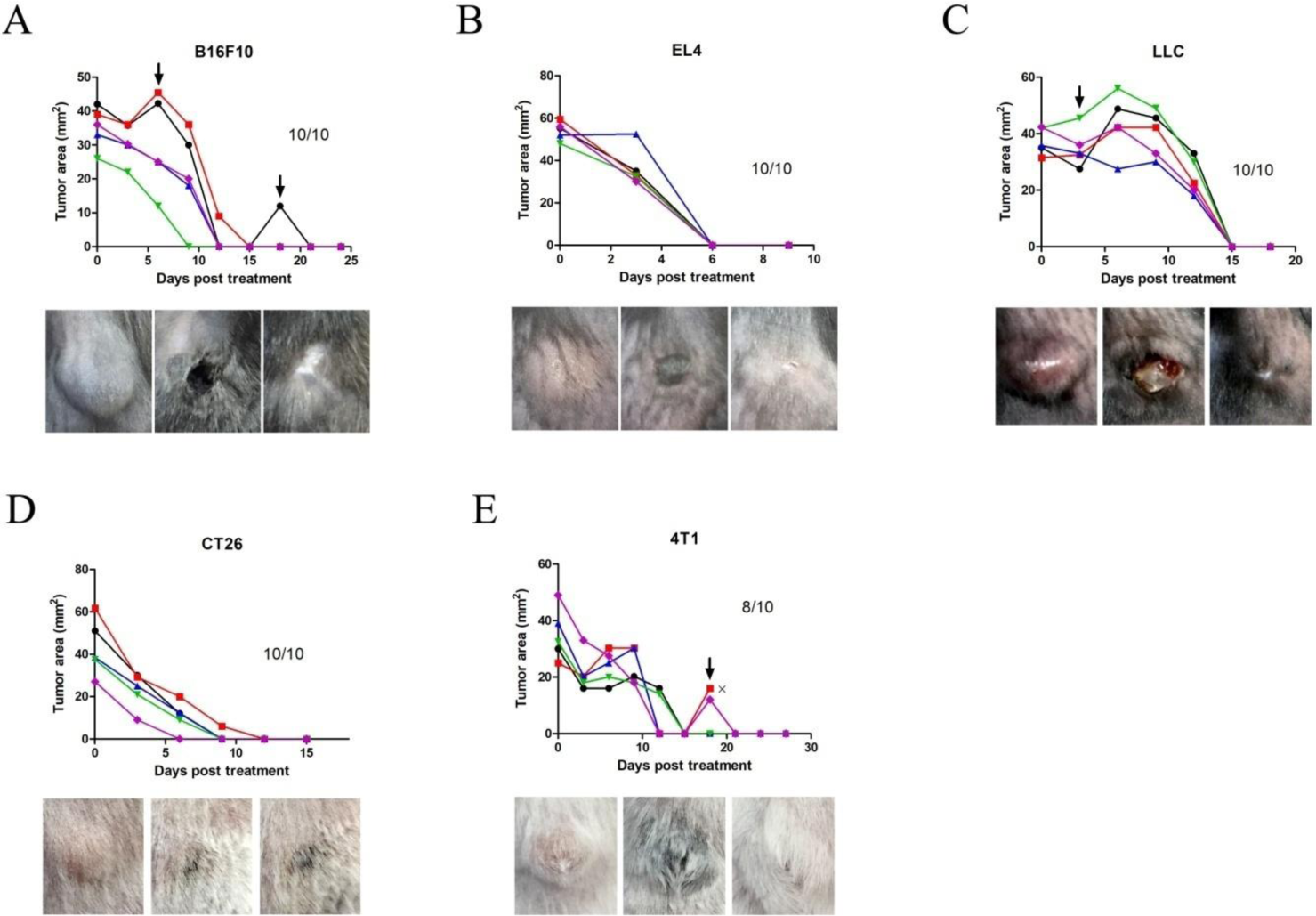
Treatment of syngeneic tumors by intratumoral injection of chitosan/IL12+GMCSF+IL2. B16F10 (A), EL4 (B), LLC (C), CT26 (D) and 4T1 (E) tumors were established by subcutaneous injection of tumor cells into the flank of syngeneic mice. Intratumoral injection of chitosan/IL12+GMCSF+IL2 was performed when the tumor diameter reached ~6 mm. Tumor growth was monitored and additional injections were carried out at the signs of tumor size increase or relapse after regression, which was indicated by arrows in the graphs. X indicated a death case. Numbers in the graphs indicated tumor cleared mice/total treated mice. Photographs indicated a representative tumor regression progress.

Next the therapeutic potential of chitosan/IL12+GMCSF+IL2 was evaluated in other murine syngeneic tumor models, including lymphoma cell (EL4), Lewis lung carcinoma (LLC), colon carcinoma cell (CT26) and breast cancer cell (4T1). Single injection of chitosan/IL12+GMCSF+IL2 led to complete regression of EL4 and CT26 tumor lesions (Fig. 4B, Fig. 4D). All LLC tumors regressed after 2 round intratumoral injections (Fig. 4C). Part of 4T1 tumors relapsed 1-2 weeks after regression. Apart from 2 mice died within 4 days post secondary injection, the remaining cleared the relapsed 4T1 tumors, generating a tumor free long-term survival (Fig. 4E). Regardless of various cancer immunograms, all tumor bearing mice were curable by intratumoral injection once the therapeutics was tolerated. In addition, an old mice harboring a spontaneous tumor at ear root was treated with chitosan/IL12+GMCSF+IL2. After injection, the tumor gradually regressed within 10 days (fig. S13). It demonstrated that this treatment, targeting the immune system, could be applied on various malignant tumors. To test the therapeutic potential in large animals, two dogs were treated by intratumoral injection of chitosan/IL12+GMCSF+IL2. CR in the dog with oral malignant melanoma and PR in the dog with mammary carcinoma were induced by a single injection, without serious adverse effects (fig. S14).

Cytokines has been used in cancer treatment clinic for decades, mostly in systemic administration. Poor therapeutic outcomes and high side effects greatly restricted their extensive application (25, 26). Local delivery of cytokines using biomaterials has been attempted in some murine tumor models, as well as synergistic effect was observed in combination usage (27–29). Because of the infinite cytokine combinations, it is unaffordable to screen cytokines with high antitumor activities using those methods. However, the lentivector based inducible system presented here resolved these problems. Induction of cytokine expression in tumor cells provides sustained local cytokine enrichment, greatly improving the lesion/serum ratio of drug concentration, which ultimately unmasks the therapeutic potential of cytokines. Albeit inferior to inducible expression, chitosan/IL12+GMCSF+IL2 treatment erased all tested murine tumors, suggesting the practicability of this screen system. It is noteworthy that in situ administration is crucial, as systemic IL2 delivery attenuate the antitumor effect of IL12+GMCSF (30). The lethal toxicity observed in mice with high tumor burden might be tolerated in human patients as the much lower tumor/body weight ratio compared to small animals. Recently, the T cell response to immune checkpoint therapy in human patients was found to be similar with that in murine tumor model (31). Considering many human cytokines have been on the market, our study in experimental mice is valuable for rapid improving immune therapeutic strategies targeting human malignant neoplasm.

## Acknowledgments

We thank L. Y. Zou and L. Fei for the assistance in mouse operation. We thank W. W. Zeng for the direction in IHC staining.

## Materials and Methods

### Cell culture

Mouse B16F10 melanoma cell line, Lewis lung carcinoma (LLC) cell line, EL4 lymphoma cell line and CT26.WT colon carcinoma were maintained in Dulbecco’s modified Eagle’s Medium (DMEM) supplemented with 10% FBS (Life Technologies), penicillin/streptomycin (Life Technologies) at 37 °C in 5% CO2 incubator. Mouse 4T1 mammary carcinoma cell line was maintained in RPMI1640 medium supplemented with 10% FBS (Life Technologies), penicillin/streptomycin (Life Technologies) at 37°C with 5% CO2. Human embryonic kidney cell line 293 and 293FT were maintained in DMEM with 10% FBS (Life Technologies), penicillin/streptomycin (Life Technologies) at 37°C in 5% CO2 incubator.

### Plasmid construction and cell transduction

The DNA sequence encoding a reverse tet transactivator protein (rtTA) was synthesized (Genscript) and subcloned into lentiviral vector pLentis-CMV-MCS-IRES-Bsd between BamHI and XhoI sites, generating the lentiviral vector pLentis-CMV-rtTA-IRES-Bsd. GFP, iRFP, IFNa, ΙΡΝγ, lymphotoxin, IL12, GMCSF, FLT3L, IL2, IL15 and IL21 coding sequences were synthesized and subcloned into lentiviral vector pLentis-TRE-MCS-PGK-PURO between BamHI and XhoI sites, generating the inducible vector pLentis-TRE-Genes-PGK-PURO. The transcription of inserted genes was controlled by the tet-responsive TRE promoter. The two subunits of lymphotoxin, Ltα and Ltβ, were jointed using a P2A sequence. The two subunits of IL12, IL 12α and IL 12β, were jointed using a T2A sequence. The cytokine expression vectors were constructed by ligated IL12, GMCSF, FLT3L or IL2 coding sequences into BamHI and XhoI sites pLentis-CMV-MCS-IRES-PURO, resulting in pLentis-CMV-IL12-IRES-PURO, pLentis-CMV-GMCSF-IRES-PURO, pLentis-CMV-FLT3L-IRES-PURO and pLentis-CMV-IL2-IRES-PURO.

The lentiviral particles were produced by cotransfection of pMD2.G, psPAX2 and lentiviral vectors into 293FT cells. B16F10 cells were firstly tranduced with pLentis-CMV-rtTA-IRES-Bsd virus and selected with 8 ug/ml blasticidin, generating B16F10-rtTA cells. Subsequently, B16F10-rtTA cells were transduced with pLentis-TRE-Genes-PGK-PURO virus and selected with 3 ug/ml puromycin, generating doxycycline (DOX) inducible cells, B16F10-rtTA (TRE-Genes). The transduced cells were selected with blasticidin and puromycin every two passages. To construct cytokine producing cells, 293 cells were transduced with pLentis-CMV-IL12-IRES-PURO, pLentis-CMV-GMCSF-IRES-PURO, pLentis-CMV-FLT3L-IRES-PURO or pLentis-CMV-IL2-IRES-PURO virus and selected using 3 ug/ml puromycin, named with 293(IL12), 293(GMCSF), 293(FLT3L) and 293(IL2).

### Fluorescence induction

B16F10-rtTA (TRE-GFP) cells were plated into a 24 well plate at 2 × 10^4^ cells/well. The GFP expression was induced by addition of 100 ng/ml dox immediately after plating. The cells were photographed using a fluorescence microscope (Olympus IX71).

10^5^ B16F10-rtTA (TRE-iRFP) cells were subcutaneously inoculated into the right flank of C57BL/6 mice. When the tumor grew to a long diameter of 10mm, the mice were fed with water containing 2 g/L dox. The fluorescence was detected using IVIS spectrum in vivo imaging system (Caliper Quantum FX).

### Mice

C57BL/6 and BALB/c mice were purchased from the animal center of the Third Military Medical University. Mice aged from 6-10 weeks were used for tumor inoculation. All animal experiments were conducted in accordance with guideline for the care and use of laboratory animals. At the signs of illness (lethargy, hunched posture, scruffy coat, social isolation, inactivity or weight loss), mice were taken out of the experiments. The investigators were not blinded in experiment conduction and data analysis. Animal experiments were conducted in at least 2 separate environments to avoid the result bias caused by microbiome.

### Assessment of the effects of induced cytokine expression on tumors

In experiments evaluating the effects of single cytokine expression on tumor growth, 105 inducible tumor cells were subcutaneously injected into the right flank of mice. In experiments assessing the effects of cytokine combinations, tumor cell mixtures containing 5×10^4^ of each inducible tumor cell line were subcutaneously injected into the right flank of mice. When tumor nodule noticeably formed, dox was continuously administered by adding 2g/L dox into drinking water. The perpendicular diameters of tumors were measured using a caliper every 3 days, and tumor area was calculated by a×b, where a was the long diameter and b was the short diameter. In experiments investigating the therapeutic effects of induced IL12+GMCSF+IL2 expression on large tumors, dox was administered after long diameter reached 10-15mm or over 15mm.

To detect the existence of immune memory2× 10^5^ parental B16F10 cells were intravenously injected or subcutaneously inoculated into the contralateral flank of cured mice 2 month after primary tumor clearance. Subcutaneous tumor formation and mice survival were recorded daily.

In experiments exploring the effects of induced IL12+GMCSF+IL2 expression on tumors at contralateral flank, mice were allocated into 3 groups based on the tumor sizes. 1, Inducible tumor cells were s.c. injected at right flank 10 days prior to 10^5^ parental tumor cells inoculation at contralateral flank. At the time of induction, the inducible tumors were larger (~10mm in diameter) and the parental tumors were smaller (~3mm in diameter). 2, Inducible tumor cells were s.c. injected at right flank 1 days prior to 10^5^ parental tumor cells inoculation at contralateral flank. At the time of induction, the inducible tumors were smaller (~5mm in diameter) and the parental tumors were larger (~15mm in diameter). 3, Inducible tumor cells were s.c. inoculated at right flank of mice. When the tumors were palpable, 10^5^ parental tumor cells were s.c. injected at contralateral flank. At the time of induction, the sizes of tumors at both flanks were comparable (~10mm in diameter).

In experiments exploring the effects of induced IL12+GMCSF+IL2 expression on i.v. infused tumor cells, mice were allocated into 2 groups based on the time of induction. Inducible tumor cells were s.c. inoculated at right flank of mice. 2× 10^5^ parental tumor cells were i.v. injected after the tumor nodules formed. Dox was administered immediately or 2 days later. Mice survival was monitored daily.

### Detection of cytokine secretion

Inducible tumor cells were plated into a 24 well plate at 5× 10^4^ cells/well in 700ul medium. 100 ng/ml dox was separately administered at 24h, 48h or 72h, all supernatants were collected at 96h after cell plating.

293 cells expressing IL12, GMCSF or IL2 were plated into a 24 well plate at 5× 10^4^ cells/well in 700ul medium. The supernatants were collected 96h later.

The concentration of IL12p70, GMCSF or IL2 in the supernatants was measured with ELISA Kits (Neobioscience), according to the manufacturer’s instructions.

### Tumor infiltrating immune cell analysis

Mice bearing tumors of ~6mm in diameter were used in this experiment. At 0 day, 1 day, 2 day and 3 day after dox administration, mice bearing inducible tumors were sacrificed. In control group, mice bearing parental tumors were sacrificed at 0 day and 3 day after dox addition. Tumor tissues were isolated with scalpel and fixed in 1% paraformaldehyde overnight. After rinsed in PBS, the samples were hydrated in 30% sucrose/PBS overnight. The tissues were mounted in OCT embedding compound and cut in 10um tissue sections using a cryostat. The sections were fixed in PBS /1% PFA, permeabilized in PBS /0.2% Tween-20 /0.3M glycine and blocked with PBS /0.2% Tween-20/5% heat-inactivated FBS/0.05% NaN3. Then the sections were incubated with 1:500-1:1000 dilution for each primary antibody in PBS/0.2% Tween-20/5% heat-inactivated FBS/0.05% NaN3. After washes, the sections were incubated with 1:500 dilution for corresponding secondary antibodies in PBS/0.2% Tween-20/5% heat-inactivated FBS/0.05% NaN3, together with trace amount of DAPI. The photographs were captured using an inverse research microscope (Nikon Ti-E).

Primary antibodies used in this study were rat anti-CD3(R&D,MAB4841-SP), rat anti-CD19(abcam, ab25232), hamster anti-CD11c(AbD Serotec, MCA1369), rat anti-F4/80(AbD Serotec, MCA497G). Secondary antibodies used in this study were Goat anti-Hamster IgG (H+L) Secondary Antibody, Alexa Fluor 647 (Invitrogen, A-21451), Donkey anti-Rat IgG (H+L) Highly Cross-Adsorbed Secondary Antibody, Alexa Fluor 488 (Invitrogen, A-21208).

### Cytokine solution production

293 cells expressing IL12, GMCSF, FLT3L or IL2 were plated into 15 cm dishes in complete DMEM medium. After the cells grew over 90% confluence, culture medium was replaced with 25 ml CDM4HEK293 medium (Hyclone) supplemented with 4 mM L-Glutamine. 96h later, the supernatants were collected and filtrated through 0.45 um filter (Millipore). Next, each cytokine supernatant was concentrated to 1 ml using Amicon Ultra-15 centrifugal filter unit (Millipore), according to the manufacturer’s instructions. The cytokine solutions were aliquoted and stored at −20 °C.

### Murine tumor treatment

10^5^ B16F10, 2×10^5^ LLC or 10^6^ EL4 cells were s.c. injected into the right flank of C57BL/6 mice. 5×10^5^ 4T1 or 5×10^5^ CT26 cells were s.c. injected into the right flank of BALB/c mice. Treatment was initiated when the tumor reached ~6mm in diameter.

In experiments using cells for treatment, 293 cells expressing IL12, GMCSF or IL2 were digested and resuspended in complete DMEM medium at a density of 1.5× 10^5^ 293-IL12 +

1.5×10^5^ 293-GMCSF + 1.5×10^5^ 293-IL2/45 ul. After addition of 1.5 ul 2M CaCl2, 45 ul 1.5% algilate (Sigma Aldrich, A0682) solution was added and mixed immediately to form hydrogel, which was slowly injected into tumors at 1.5 ul/mm2 using a 29G insulin syringe (Becton).

In experiments using cytokine solution for treatment, 45 ul solution containing 15 ul of each cytokine were mixed with 45 ul 3% chitosan (Chitosan glutamate, Protosan G 213, NovaMatrix) solution. Then the chitosan/cytokine solution was slowly injected into tumors at 1.5 ul/mm2 using a 29G insulin syringe.

At the signs of disease progress, e.g. tumor size increase or new nodule appearance, additional injections were conducted until the tumor reached 12 mm in diameter.

A 50-week BALB/c mice bearing a 10mm× 10mm spontaneous tumor at ear root was intratumorally injected with 100 ul chitosan/IL12+GMCSF+IL2.

### Dog tumor treatment

Two dogs, diagnosed with advanced stage cancer, were treated with chitosan/IL12+GMCSF+IL2 at animal hospital of beijing chaoyang district animal disease control center. Informed consent signature by dog owners was required before treatment. Canine IL12, GMCSF and IL2 (Novas Biologicals) was reconstituted to 150 ng/ul. The cytokine solution for injection was prepared by mixing IL12, GMCSF, IL2 and 3% chitosan at a volume ratio of 1:1:1:3. The dose of injection was 1 ul cytokine solution per mm^2^ tumor area. Solutions were slowly injected into nodules using a 29G insulin syringe and the status of dogs was closely monitored. Treatment response was defined as follows: complete response (CR), all tumors disappearance; partial response (PR), at least 30% reduction in the sum of long tumor diameter; progressive disease (PD), at least 20% increase in the sum of long tumor diameter.

### Statistics

Statistical analysis was carried out in GraphPad Prism 5 software. Survival curves were analyzed with the log-rank (Mantel-Cox) test. *p*<0.05 was considered as significant, \**p*<0.05,\*\**p*<0.01,\*\*\**p*<0.001.

---

**fig. S1.**
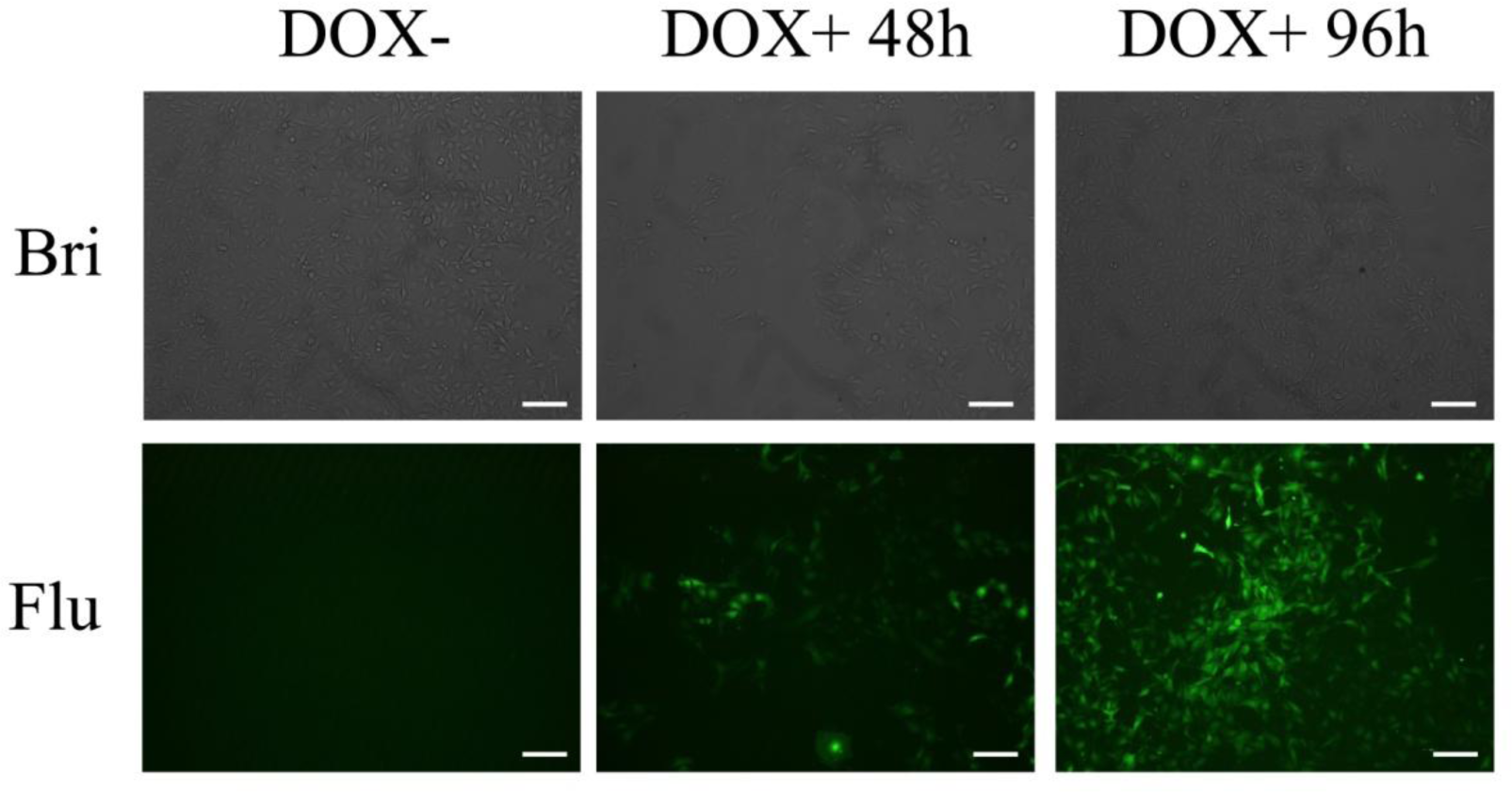
*In vitro* assessment of the inducible system in B16F10 cells. B16F10-rtTA(TRE-GFP) cells were plated into 24 well plate at 2×10^4^ cells/well. 100 ng/ml dox was added just after plating, and wells without dox addition served as control. Fluorescence was photographed at 48h and 96h after induction. Scale bars, 50 um.

**fig. S2.**
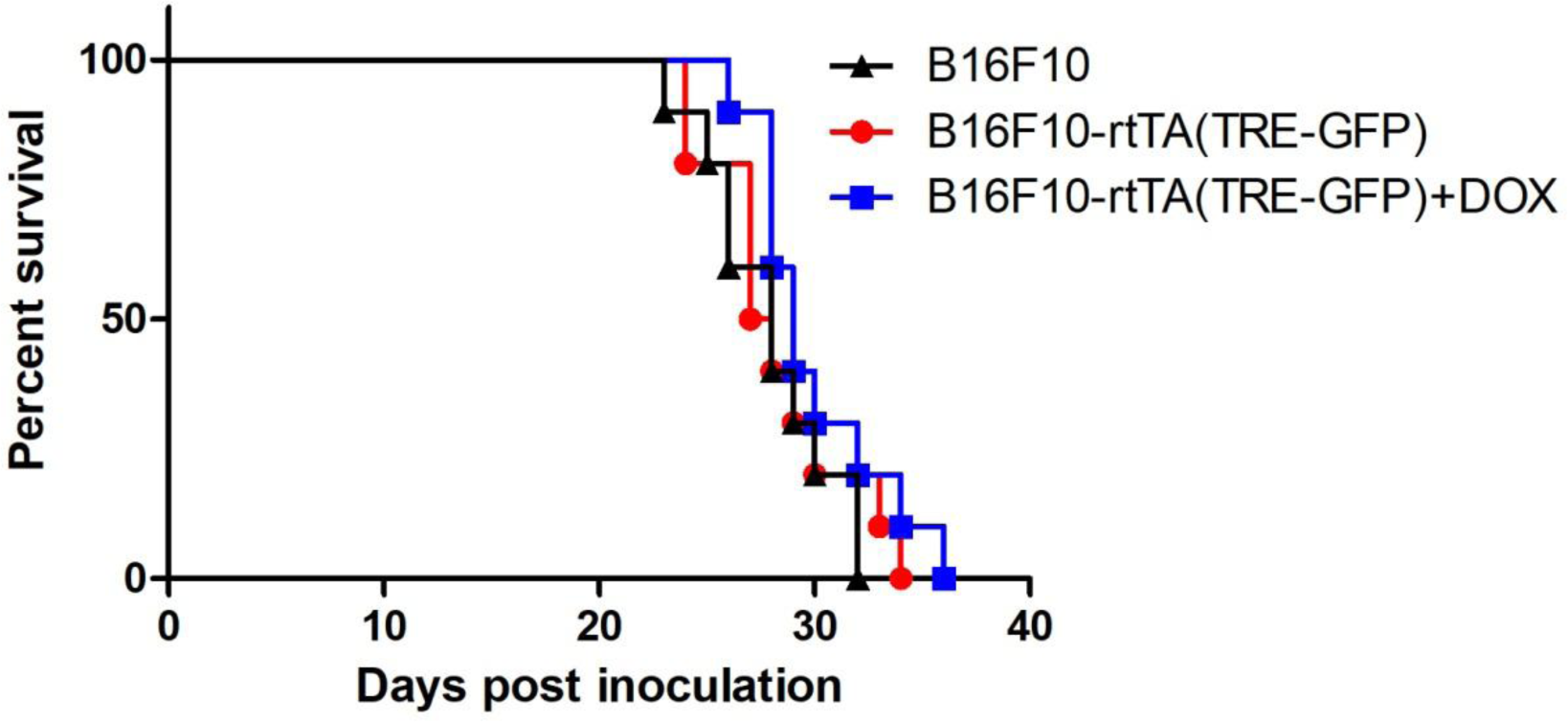
The influence of dox and lentiviral transduction to the survival of tumor bearing mice. Mice were allocated into three groups with different treatment, (1) subcutaneously inoculated with 10^5^ parental B16F10 cells, (2) subcutaneously inoculated with 10^5^ B16F10-rtTA(TRE-GFP) cells and (3) subcutaneously inoculated with 10^5^ B16F10-rtTA(TRE-GFP) cells with dox supplement in drinking water. The survival of tumor bearing mice were recorded. n=10.

**fig. S3.**
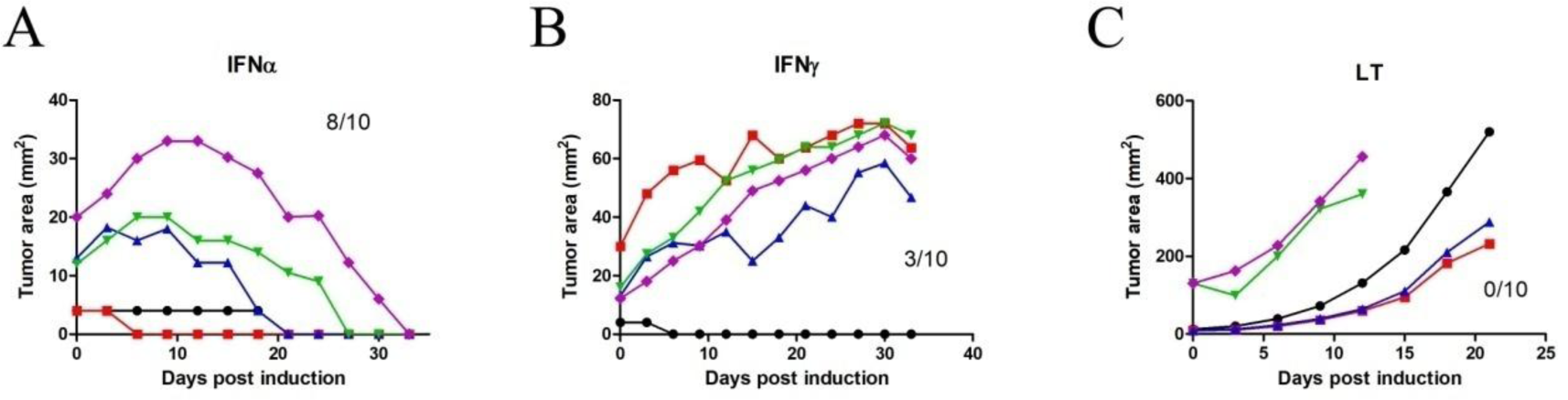
Effects of induced cytokine expression on the growth of established tumors. (A) 10^5^ B16F10-rtTA(TRE-IFNα), (B) B16F10-rtTA(TRE-IFNγ) or (C) B16F10-rtTA(TRE-LT) cells were subcutaneously injected into the flank of mice. Dox was administered after tumor nodule formation. The tumor size was monitored every 3 days after induction. Each curve represented the tumor growth of an individual mouse after dox administration. Numbers in the graphs indicated tumor cleared mice/total dox treated mice.

**fig. S4.**
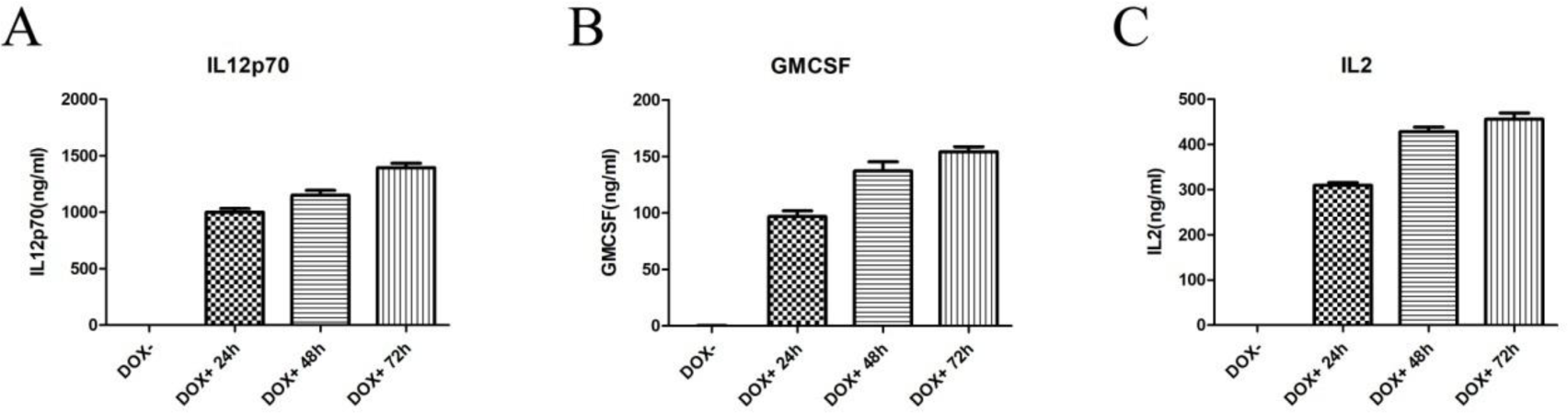
The expression level of IL12, GMCSF and IL2 in inducible B16F10 cells after induction. (A) B16F10-rtTA(TRE-IL 12), (B) B16F10-rtTA(TRE-GMCSF) or (C) B16F10-rtTA(TRE-IL2) cells were plated into 24 well plate at 5 × 10^4^ cells/well. 100 ng/ml dox was separately added at 24h, 48h or 72h post plating. Wells without dox addition served as control. At 96h, all supernatants were collected and subjected to ELISA analysis of secreted cytokine concentration. n=3.

**fig. S5.**
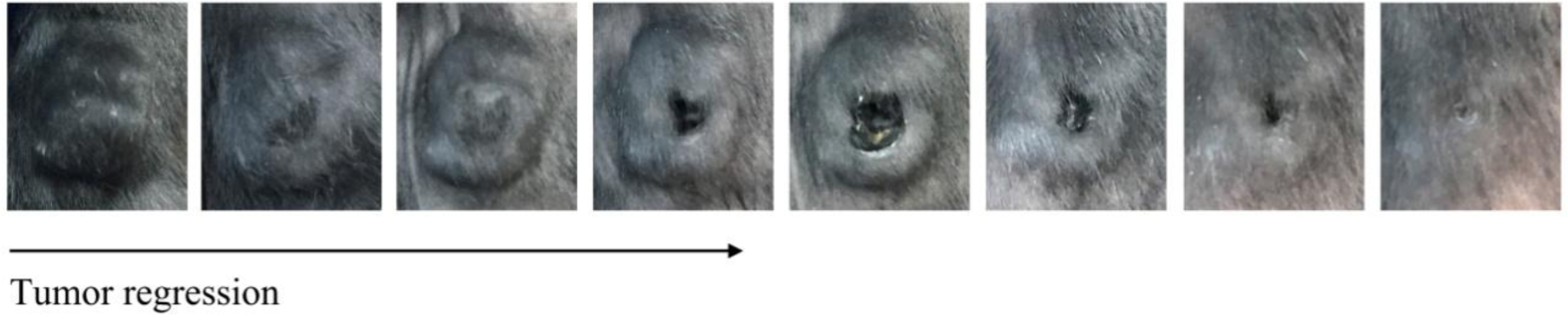
Representative photos of large tumor regression progress. 5 × 10^4^ B16F10-rtTA(TRE-IL12)+ 5 × B16F10-rtTA(TRE-IL 12)+ 5 × 10^4^ B16F10-rtTA(TRE-IL 12) cells were subcutaneously injected into the flank of mice. 2 g/L dox was administered in drinking water after the tumor was over 15mm in diameter (15.5mm × 10mm). After induction, the tumor gradually ulcered and regressed.

**fig. S6.**
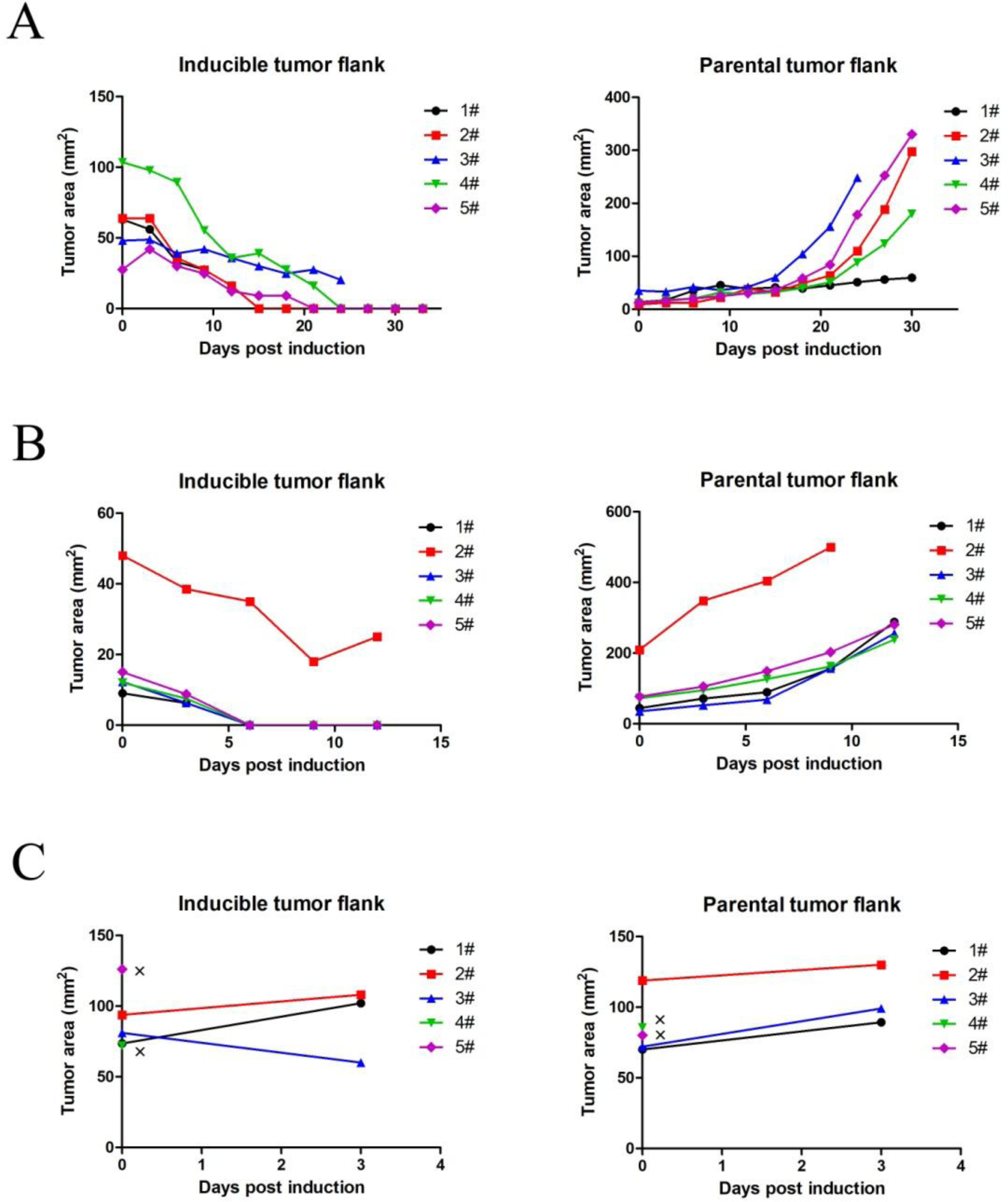
Tumor growth of mice bearing bilateral tumors. At different times post subcutaneous inoculation of inducible tumor cells (IL12+GMCSF+IL2) into the flank of C57BL/6 mice, parental tumor cells were subcutaneously injected into the contralateral side. Dox induction was initiated when the size of bilateral tumors at different ratio: (A) inducible tumor larger, (B) parental tumor larger, or (C) sizes of both tumors were comparable. The growth of both tumors was monitored every 3 days after induction. X indicated a death case within 6 days after induction. n=5.

**fig. S7.**
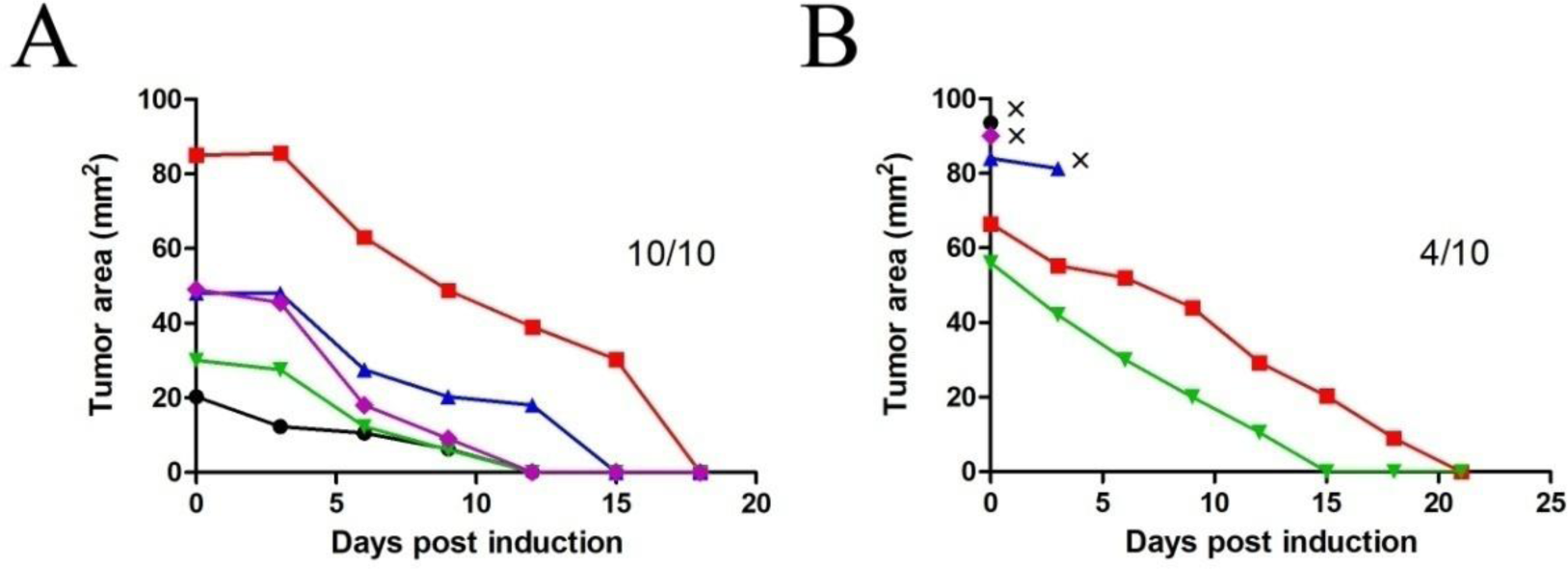
Subcutaneous tumor growth of mice receiving intravenous tumor injection. Parental tumor cells were intravenously injected into mice bearing subcutaneously established inducible tumors (IL12+GMCSF+IL2). Dox induction was initiated just after i.v. injection (A) or 2 days later (B). The sizes of subcutaneous tumors were measured every 3 days after induction. X indicated a death case within 4 days after induction. n=5.

**fig. S8.**
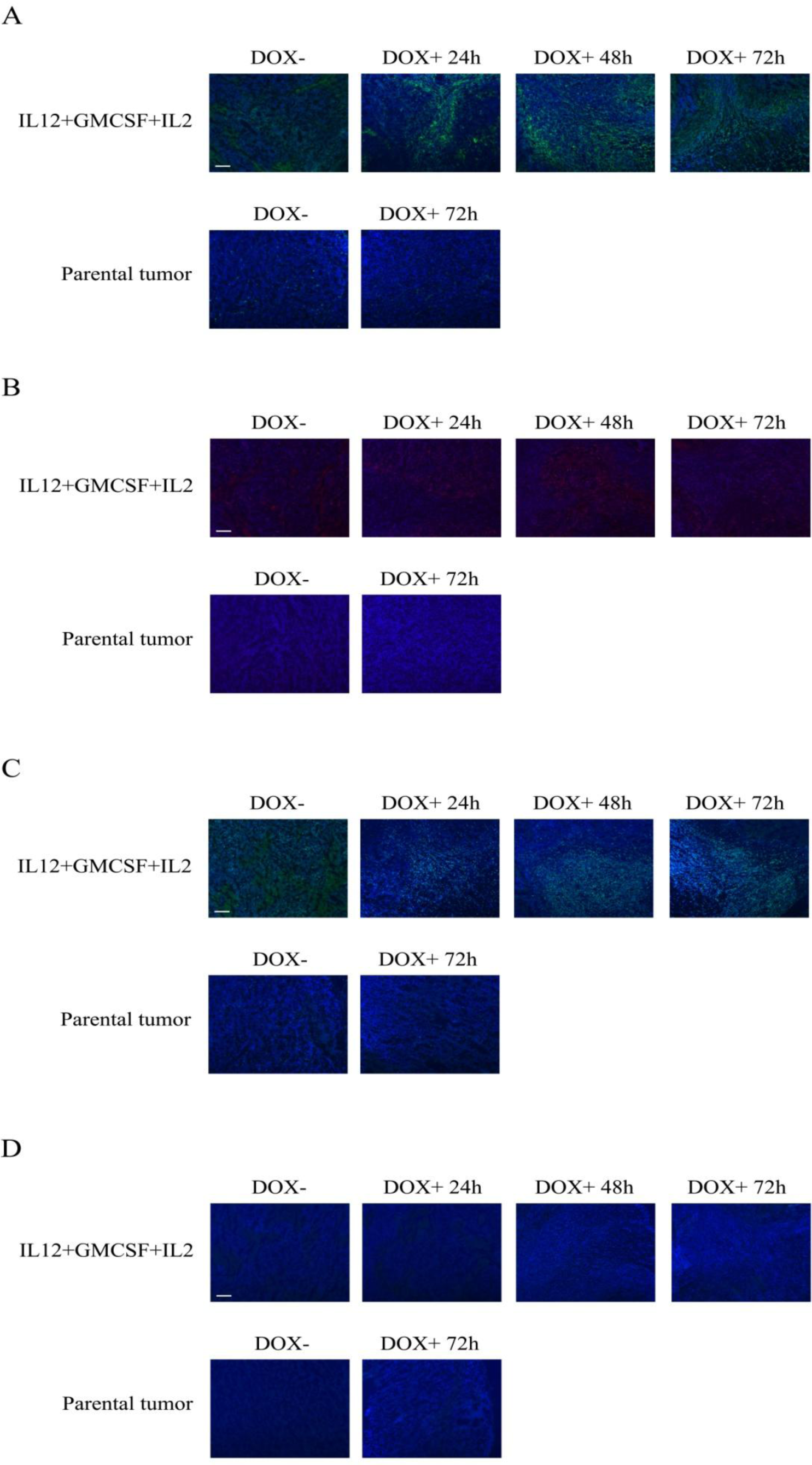
Immunohistochemistry analysis of tumor infiltrating immune cells after induciton. Mice bearing tumors of ~6mm in diameter were administered with dox in drinking water. In the group of bearing parental tumor, mice were sacrificed just prior dox administration or 3 days later. In the group bearing inducible tumor (IL12+GMCSF+IL2), mice were sacrificed just prior dox administration or 1, 2, 3 days post induction. Tumor tissues were collected and subjected to IHC staining of F4/80 (A), CD11c (B), CD3 (C) and CD19 (D). Scale bars, 100 um.

**fig. S9.**
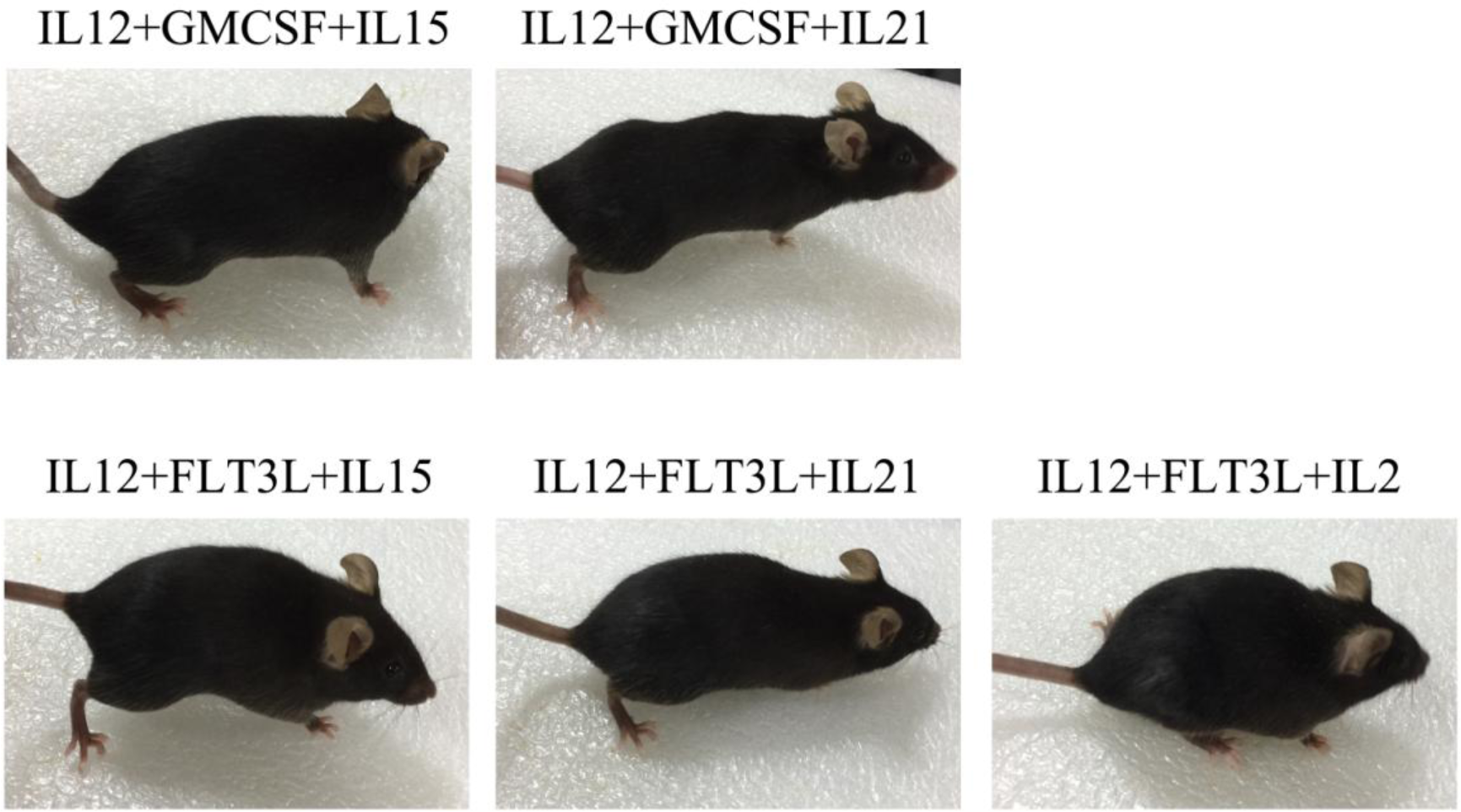
Representative vitiligo appearance in mice with tumor regression after dox induction, including 5 inducible tumor cell combinations: IL12+GMCSF+IL15, IL12+GMCSF+IL21, IL12+FLT3L+IL15, IL 12+FLT3L+IL21, IL12+FLT3L+IL2.

**fig. S10.**
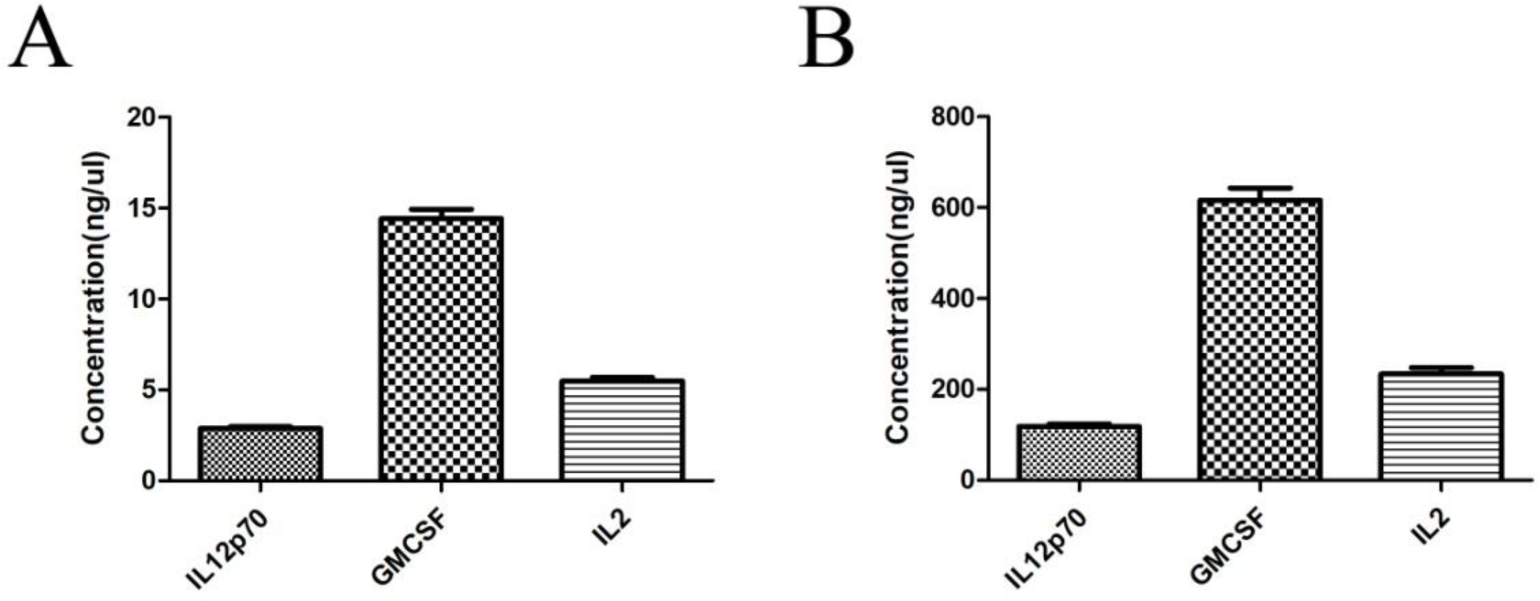
Concentration of IL12, GMCSF and IL2 in the supernatants from transduced 293 cells and ultrafiltrated cytokine solutions. (A) 293(IL12), 293(GMCSF) and 293(IL2) cells were separately plated into 24 well plate at 5 ×10^4^ cells/well. 96h later, the supernatants were collected and subjected to ELISA analysis. (B) The supernatants from 15cm dish culture of 293(IL12), 293(GMCSF) or 293(IL2) cells were concentrated by ultrafiltration. Cytokines in the concentrated solutions were measured using ELISA analysis. n=3.

**fig. S11.**
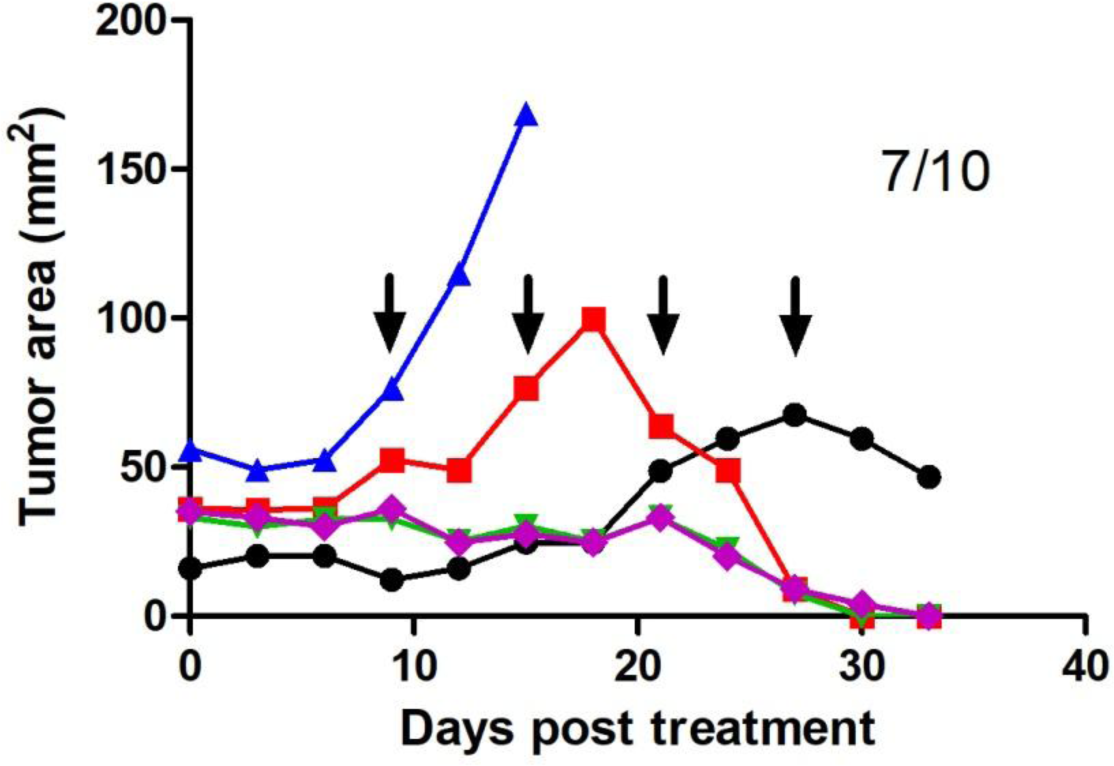
Treatment of B16F10 tumors using agilate hydrogel/293(IL12)+293(GMCSF)+293(IL2) cells. B16F10 cells were subcutaneously inoculated into the flank of mice. As tumor reached ~6mm in diameter, agilate hydrogel containing 293(IL12), 293(GMCSF) and 293(IL2) cells were injected into tumors. Additional injections were performed at the sign of tumor size increase, which was indicated by arrows in the graph. Tumor growth was monitored every 3 days after treatment. Numbers in the graph indicated tumor cleared mice/total treated mice.

**fig. S12.**
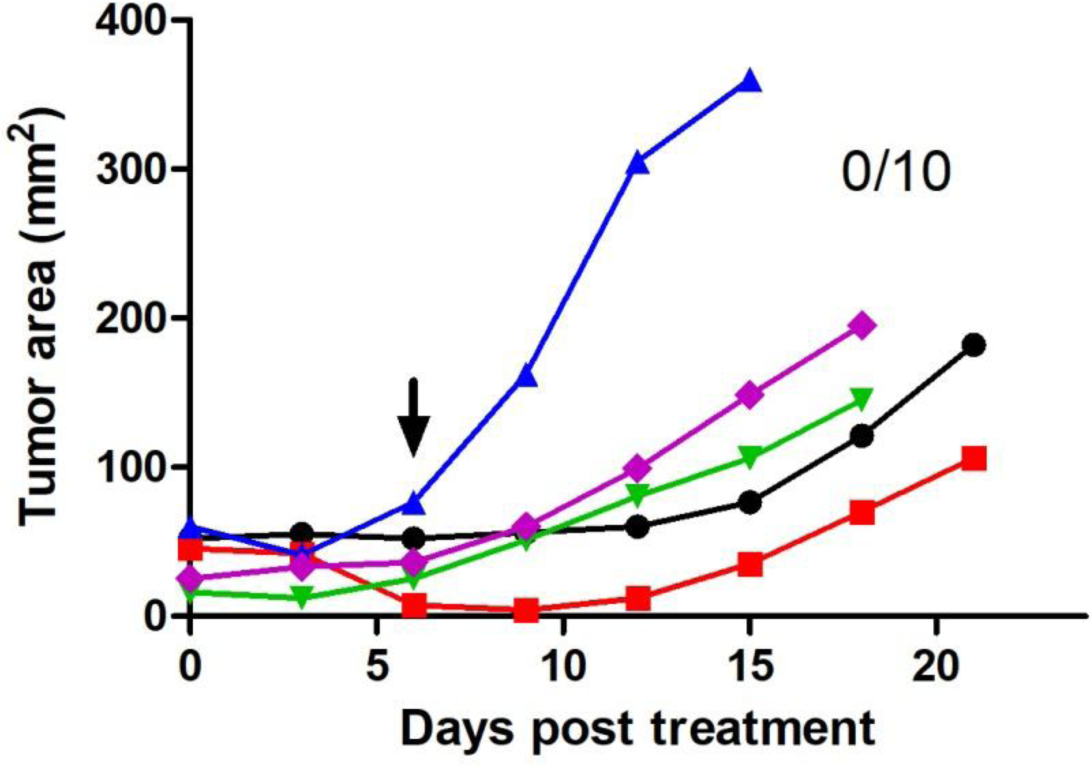
Treatment of B16F10 tumors using chitosan/IL12+FLT3L+IL2. 10^5^ B16F10 cells were subcutaneously inoculated into the flank of mice. As tumor reached ~6mm in diameter, chitosan/IL12+FLT3L+IL2 were injected into tumors. Additional injections were performed at the sign of tumor size increase, which was indicated by arrows in the graph. Tumor growth was monitored every 3 days after treatment. Numbers in the graph indicated tumor cleared mice/total treated mice.

**fig. S13.**
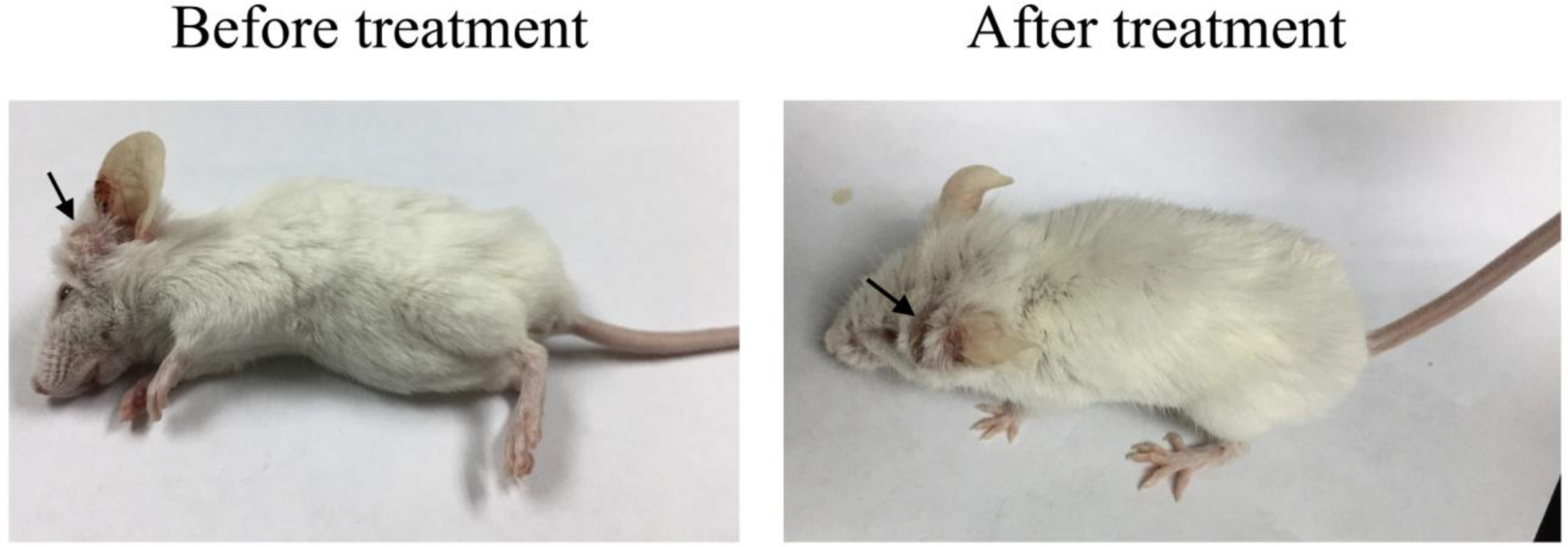
Treatment of a mouse bearing spontaneous tumor using chitosan/IL12+GMCSF+IL2. A 10mm spontaneous tumor was observed at the ear root of a BALB/c mouse. Injection of chitosan/IL12+GMCSF+IL2 led to a complete regression of this tumor. Arrows indicated the tumor site.

**fig. S14.**
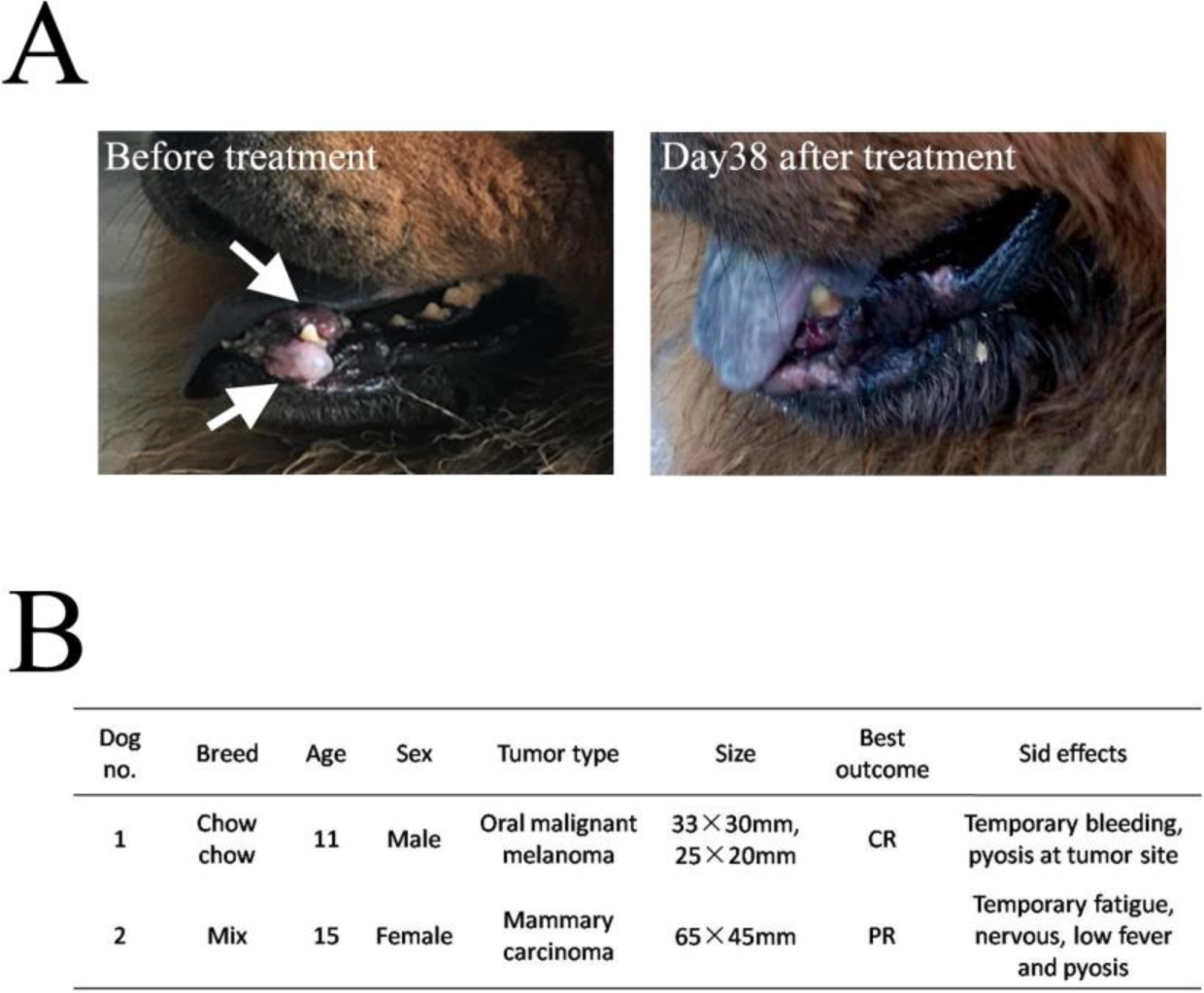
Treatment of dogs diagnosed with malignant tumor using chitosan/IL12+GMCSF+IL2. (A) Photographs of dog 1 with oral malignant melanoma. (B) Overview of treatment information.

